# Expression, Purification, and Functional Reconstitution of ^19^F-labeled Cytochrome b5 in Peptide Nanodiscs for NMR Studies

**DOI:** 10.1101/844381

**Authors:** Jia Bai, Jian Wang, Thirupathi Ravula, Sang-Choul Im, G. M. Anantharamaiah, Lucy Waskell, Ayyalusamy Ramamoorthy

## Abstract

Microsomal cytochrome b5 (cytb5) is a membrane-bound protein capable of donating the second electron to cytochrome P450s (CYP P450s) in the CYP P450s monooxygenase reactions. Recent studies have demonstrated the importance of the transmembrane domain of cytb5 in the interaction with cytP450 by stabilizing its monomeric structure. While recent NMR studies have provided high-resolution insights into the structural interactions between the soluble domains of ~16-kDa cytb5 and ~57-kDa cytP450 in a membrane environment, there is need for studies to probe the residues in the transmembrane region as well as to obtain intermolecular distance constraints to better understand the very large size cytb5-cytP450 complex structure in a near native membrane environment. In this study, we report the expression, purification, functional reconstitution of ^19^F-labeled full-length rabbit cytb5 in peptide based nanodiscs for structural studies using NMR spectroscopy. Size exclusion chromatography, dynamic light scattering, transmission electron microscopy, and NMR experiments show a stable reconstitution of cytb5 in 4F peptide-based lipid-nanodiscs. Our results demonstrate that the use of peptide-nanodiscs containing cytb5 ^19^F-labeled with 5-fluorotryptophan (5FW) enables, for the first time, the detection of residues from the transmembrane domain in different lipid compositions by solution ^19^F NMR experiments. ^19^F NMR results revealing the interaction of the transmembrane domain of cytb5 with the full-length rabbit Cytochrome P450 2B4 (CYP2B4) are also presented. We expect the results presented in this study to be useful to devise approaches to probe the structure, dynamics and functional roles of transmembrane domains of a membrane protein, and also to measure intermolecular ^19^F-^19^F distance constraints to determine the structural interactions between the transmembrane domains.

## INTRODUCTION

Despite the recent advances in high-resolution structural studies of membrane proteins, obtaining their high-resolution dynamics continues to be a major challenge. Although both solution and solid-state NMR spectroscopy can be used to characterize the dynamics of membrane proteins, low sensitivity and resolution are the key limiting factors of NMR techniques for studies on challenging membrane proteins. One of the strategies used to overcome these challenges is to selectively label the protein with ^19^F nuclei as ^19^F NMR experiments can probe the local structure of a protein as well as to measure ^19^F-^19^F distance constraints for high-resolution structural studies[1–5]. In this study, we extend this approach to study a membrane-bound cytochrome b5 (cytb5).

Microsomal cytb5 is a 134–residue electron transport hemeprotein primarily associated with the endoplasmic reticulum of eukaryotic cells, and plays a vital role in the enzymatic function of cytochrome P450s[6–8]. Cytb5 can donate the second of the two required electrons to cytochrome P450s (CYP P450s), which can catalyze the metabolism of endogenous and exogenous substrates including fatty acids, sterols and over 50% clinical drugs[8–11]. Therefore, it is very important to measure the structural and dynamical interactions between the full-length membrane-bound cytb5 and CYP P450s to better understand the catalytic mechanism of cytP450.

Cytb5 is comprised of three distinct regions: a heme containing soluble N-terminus (residue 1-89), a hydrophobic transmembrane C-terminus (residue 105-134), and a 15-residue flexible linker (residue 90-104) connecting the soluble and the transmembrane domains[6, 12–15]. Several studies have shown that the soluble domain of cytb5, lacking the transmembrane domain, aggregates to form oligomers and they do not interact with cytP450 in a physiologically-relevant manner[6, 16–18]. Moreover, our previous studies have demonstrated the significance of the lipid membrane in the structural interactions between cytb5 and CYP P450[19–22]. Therefore, to better understand the role of the transmembrane domain of cytb5, it is important to investigate the structure and dynamics of the full-length cytb5 in a membrane environment. However, the membrane-bound full-length cytb5 poses tremendous challenges for atomic-level structural studies by X-ray crystallography or solution-state nuclear magnetic resonance (NMR) spectroscopy[19–22]. The transmembrane residues of cytb5 are invisible to solution NMR spectroscopy[21], and only solid-state NMR studies are capable of probing the structure and dynamics of the transmembrane helix[22, 23].

Tryptophan is abundant in membrane proteins, mostly located at the lipid-water interface, where it is thought to play a significant role as a membrane anchor [24–27]. Computational and experimental studies report that tryptophan residues of a membrane protein contribute to the stability and orientation of transmembrane domains, and play a significant role in membrane deformations[24, 27–31]. Cytb5 has 4 tryptophan residues: W27 is located in the N-terminus while the other three Trp residues are located in the transmembrane domain. In this study, the full-length cytb5 was labeled with 5-fluorotryptophan (5FW), and we demonstrate the use of ^19^F NMR experiments to probe the transmembrane domain of ^19^F-labeled cytb5 by solution NMR experiments. Recently lipid bilayer nanodiscs have been successfully used in the structural studies of membrane proteins [21, 32–35]. Here we used peptide-based lipid nanodiscs to reconstitute cytb5 or cyb5-cytP450 2B4 (CYP 2B4) complex that exhibited stability for more than a week at room temperature[21, 36]. These nanodiscs exhibited a size ~8 nm diameter, and their fast tumbling nature allowed the detection of transmembrane residues of membrane-bound cytb5 by solution NMR.

## Materials and Methods

### Materials

Potassium phosphate monobasic and dibasic, glycerol, chloroform and 3,5-di-tert-butyl-4-hydroxytouene (BHT), 5-fluoroindole (98%) were purchased from Sigma-Aldrich. 1-palmitoyl-2-oleoyl-sn-glycero-3-phosphocholine (POPC), 1-palmitoyl-2-oleoyl-sn-glycero-3-phospho-L-serine (POPS), 1,2-dimyristoyl-sn-glycerol-3-phosphocholine (DMPC), 1-palmitoyl-2-oleoyl-sn-glycero-3-phosphoethanolamine (POPE), L-a-phosphatidylinositol (sodium salt) (PI), sphingomyelin (SM), and cholesterol (Chol) were purchased from Avanti Polar Lipids Inc.

### Protein expression and purification

Plasmids containing the genes of cytb5 were transformed into *Escherichia coli* strain C41. 5-fluorotryptophan (5FW) labeling was obtained by adding 5-fluoroindole (5FI). A single colony was picked from the LB plate and transferred into 100 ml of LB medium containing 100 μg/ml carbenicillin. It was incubated overnight at 30 °C under shaking at 200 rpm. 10 ml of the overnight culture was centrifuged, decanted the supernatant, and transferred the pellets into 100 ml TB medium supplemented with a final concentration of 100 μg/ml carbenicillin. The culture was incubated at 35 °C under shaking at 180 rpm. After 2 hours, the above culture was centrifuged, decanted the supernatant, and transferred the pellets into 1 L M9 medium supplemented with 0.1 g of each of the amino acids (except tryptophan). The 5-fluorotryptophan (5FW) labeling of cytb5 was obtained by adding 5-fluoroindole 10 minutes before adding IPTG[37]. The other expression conditions and purification procedures were similar as reported previously[12]. The sample purity is verified by running SDS-PAGE gel (Figure 1B).

**Figure 1.**
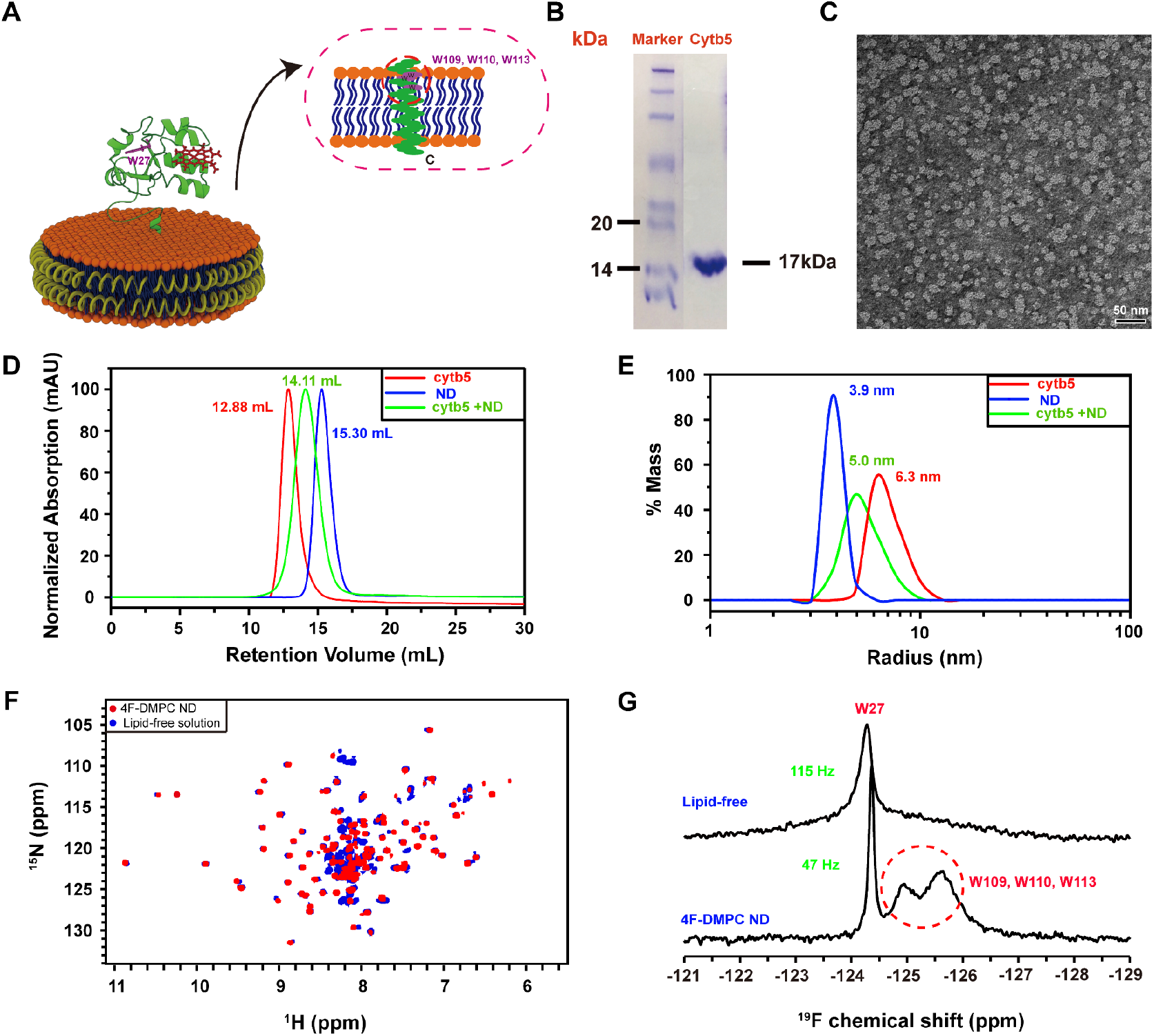
Reconstitution of cytochrome b5 in peptide-based lipid-nanodiscs. (A) Ribbon diagrams of cytochrome b5 in a nanodisc indicating 5F-tryptophan-labeled sites. (B) SDS-PAGE gel showing the purity 5FW-labeled cytb5. (C) TEM image of 4F-DMPC nanodiscs with a diameter of ~10 nm. (D) SEC elution profiles and (E) DLS profiles of cytb5 alone in buffer (red), empty 4F-DMPC nanodiscs (blue), and 4F-DMPC nanodiscs containing cytb5 (green). (F) 2D ^1^H-^15^N HSQC spectra of ^15^N-labeled cytb5 alone in buffer (blue) and reconstituted in the 4F-DMPC nanodiscs (red). (G) 1D ^19^F NMR spectra of 5F-tryptophan-labeled cytb5 alone in buffer (top trace) and reconstituted in the 4F-DMPC nanodiscs (bottom trace) at 298K.

#### Preparation of lipid-nanodiscs

Lipid powders (except DMPC) were dissolved in HPLC-grade chloroform to make a stock solution at 20 mg/mL. DMPC powder was suspended in buffer A (40 mM potassium phosphate, pH 7.4) to make a stock solution at 20 mg/mL. The 4F peptide was dissolved in buffer A to make a stock solution at 10 mg/mL. Nanodiscs were prepared as narrated below.

*Preparation of 4F-DMPC nanodiscs:* DMPC and 4F stock solution were mixed together at a peptide:lipid ratio of 1:1.5 w/w, then incubated at 37°C, 180 rpm overnight. *Preparation of 4F-POPC (or 4F-POPS, 4F-POPC/PS (8:2 molar ratio)) nanodiscs:* An aliquot of the dissolved POPC (or POPS, or POPC/PS *(8:2 molar ratio)*, 8/2, w/w) was transferred to an Eppendorf tubes, and dried under a gentle flow of nitrogen gas, followed by a 5-10 minutes drying in vacuum. The resultant lipid film was hydrated with buffer A, and mixed with 4F peptide solution at 1:1 w/w peptide:lipid. The mixture underwent 3-5 freeze-thaw cycles between −80 °C and room temperature until it became clear, then it was placed in a shaker at room temperature, 180 rpm overnight.

*Preparation of 4F-Endoplasmic Reticulum (ER) lipids-nanodiscs:* Lipids for a biomimetic ER was mixed with the following molar ratios: 58% POPC, 7% POPS, 20% POPE, 7% POPI, 4% SM, 4% chol. The mixture was dried under nitrogen gas, followed by a 5-10 minutes drying in vacuum. The resultant lipid film was hydrated with buffer A, and then mixed with 4F peptide solution at a peptide:lipid ratio of 1:0.75 w/w. The mixture underwent three to five freeze-thaw cycles between −80 °C and room temperature, then was incubated at room temperature, 180 rpm overnight. The nanodiscs were purified and characterized using size exclusion chromatography (SEC). A Superdex 200 Increase 300/10 GL column was operated on an AKTA purifier (GE healthcare, Freiburg, Germany). The radius of nanodiscs was characterized by dynamic light scattering (DLS) experiments using Wyatt Technology® DynaPro® NanoStar®. The scattering curves were fitted by the isotropic sphere model.

#### Preparation of nanodiscs containing cytb5 or the cytb5-CYP2B4 complex

To reconstitute cytb5 into nanodiscs, Cytb5 was mixed with the aforementioned purified nanodiscs at a molar ratio of 1:1.2 cytb5:nanodiscs. The mixture was then incubated at room temperature (or 37 °C), 180 rpm overnight. SEC was subsequently used for the purification of nanodiscs containing cytb5 from free cytb5 and empty nanodiscs. Fractions were collected and concentrated for NMR experiments. To reconstitute the cytb5-CYP2B4 complex into nanodiscs, purified cytb5 incorporated into nanodiscs was mixed with unlabeled full-length CYP2B4 at a molar ratio of 1:1.2 cytb5: CYP2B4. The mixture was then incubated at 37 °C, 180 rpm overnight. The cytb5-CYP2B4 complex incorporated into nanodiscs were purified by SEC, and concentrated for use in the NMR experiments.

### NMR experiments

NMR spectra were acquired on a Bruker 500 MHz NMR spectrometer equipped with a ^19^F/^1^H/^13^C/^15^N probe. 40 mM potassium phosphate was used as the NMR buffer. Protein was dissolved in 90% NMR buffer and 10% D2O, or reconstituted in 4F-based nanodiscs as described above. 5FW-labeled cytb5 (0.3 mM) was used to record ^19^F NMR spectra. ^19^F NMR spectra were acquired with 18.8 kHz spectral width, 16 K complex points, and a recycle delay of 2 s. Proton decoupling was applied to obtain all 19F NMR spectra presented in this study.

## Results and Discussion

### ^19^F solution NMR detects the resonances from the transmembrane domain

There are four tryptophans and three phenylalanines in cytb5. ^19^F-labeled cytb5 samples were obtained by using tryptophan with 5-fluoro-tryptophan (5FW) and phenylalanine with m-fluoro-phenylalanine. This procedure worked out well for the production of ^19^F-trptophan-labeled cytb5, but not for ^19^F-phenylalanine labeling. Hence, we only used ^19^F-trptohpan-labeled cytb5 in this study.

Cytb5 has four tryptophan (W27, W109, W110, W113), one located in the soluble domain and the other three are located in the transmembrane domain (Figure 1A). We labeled the tryptophans with 5-fluorotryptophan (5FW), and reconstituted the 5FW-labeled cytb5 in 4F-DMPC nanodiscs. The size of the nanodiscs was characterized by TEM (Figure 1C) and DLS (Figure 1E). The SEC profiles (Figure 1D) showed that cytb5 is successfully reconstituted in 4F-DMPC nanodiscs. ^19^F NMR spectra of 5FW-labeled cytb5 in the lipid-free solution and solution containing 4F-DMPC nanodiscs were acquired (Figure 1G). In the lipid-free solution, only a single narrow peak is observed, and a very low intensity broad peak can also be seen in the spectrum. On the other, in the solution containing 4F-DMPC nanodiscs, three resolved ^19^F peaks are observed. The narrowest peak is assigned to W27 residue located in the mobile soluble domain of the protein, whereas the two broad peaks marked with a red circle are assigned to W109, W110 and W113 (Figure 1G) that are located in the transmembrane domain of the protein. The W27 peak shows a line width of ~122 Hz in lipid-free solution, whereas it is ~49 Hz in 4F-DMPC nanodiscs, suggesting that cytb5 is aggregated in solution while it is a monomer in nanodiscs. On the other hand, the transmembrane Trp residues (W109, W110 and W113) are invisible in 2D ^1^H/^15^N HSQC NMR spectra of U-^15^N-cytb5 even when reconstituted in nanodiscs (Figure 1F), which is in agreement with our previous studies[21]. ^19^F NMR spectra of cytochrome b5 in lipid-free solution or reconstituted in nanodiscs were also acquired by varying the temperature of the sample (Figure 2). In the lipid-free solution, a single peak was observed regardless of the sample temperature (Figure 2A). However, in the 4F-DMPC nanodiscs, increasing sample temperature revealed four resolved ^19^F peaks due to the increasing tumbling rate of the cytb5-nanodiscs as well as due to the transition from the gel to liquid-crystalline phase (Figure 2B).

**Figure 2.**
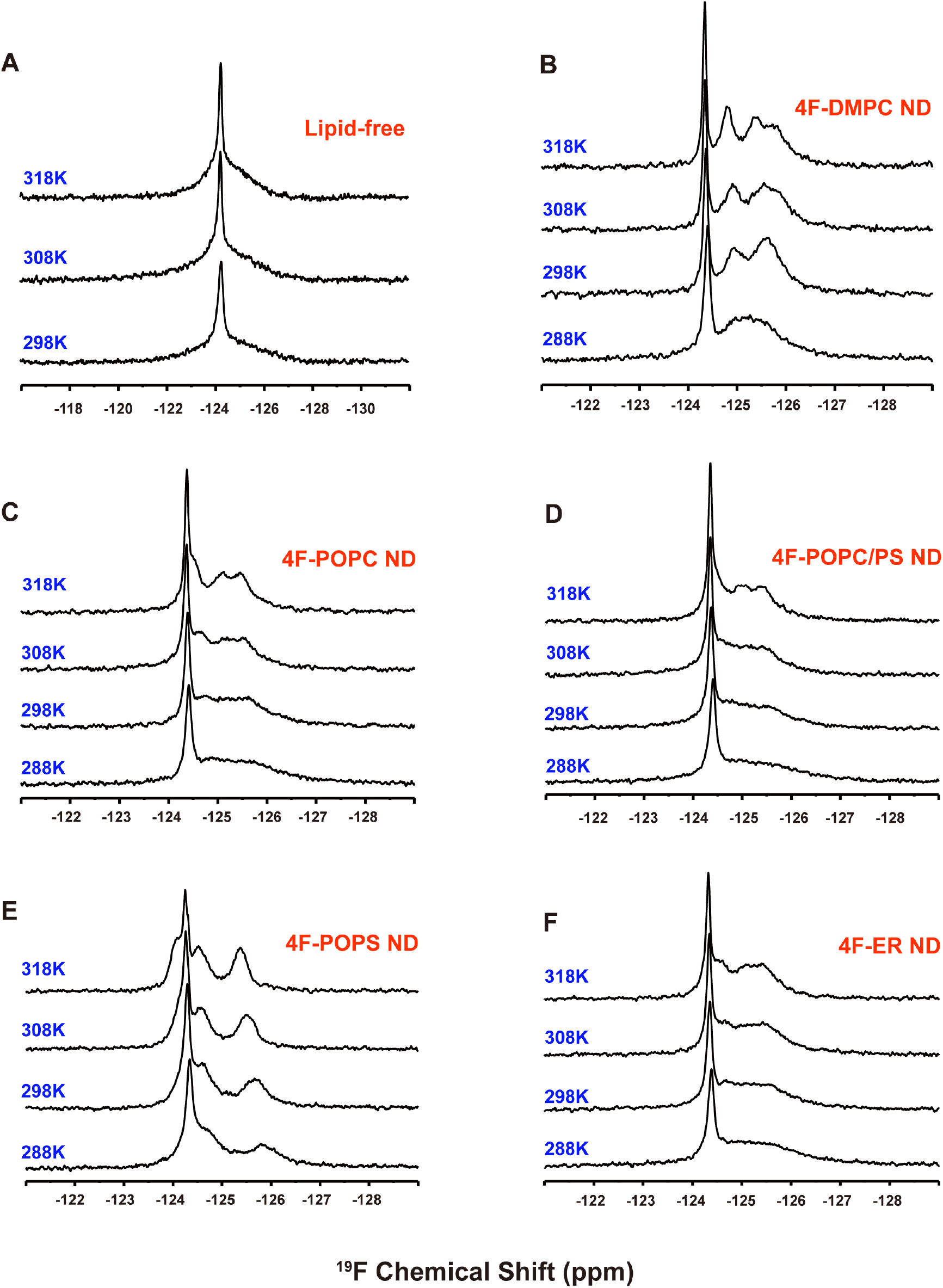
^19^F NMR spectra of 5FW-labeled cytb5 reconstituted in nanodiscs composed with different types of lipids at different temperatures. cytb5 in buffer (A) and reconstituted in the (B) 4F-DMPC, (C) 4F-POPC, (D) 4F-POPC/PS, (E) 4F-POPS, (F) 4F-ER nanodiscs at the indicated temperatures.

### Solvent PRE

Solvent PRE measurements were used to detect the solvent accessible regions of the tryptophan residues. We used Gd-DTPA-BMA as the paramagnetic cosolvent in the 4F-nanodiscs containing ^19^F-labeled cytb5, and measured the solvent paramagnetic relaxation enhancement (sPRE); and the resultant ^19^F NMR spectra are shown in Figure 3. With the increasing concentration of Gd-DTPA-BMA, a small reduction in signal intensity is observed for W27, with a relative peak height of 84% at the 5 mM Gd-DTPA-BMA. However, for the other tryptophan residues, no observable reduction in signal intensity is observed even in presence of 10 mM Gd-DTPA-BMA, (Figure 3C). These observations suggest that W27 is exposed to the solution, whereas the other tryptophans are buried in the lipid bilayer.

**Figure 3.**
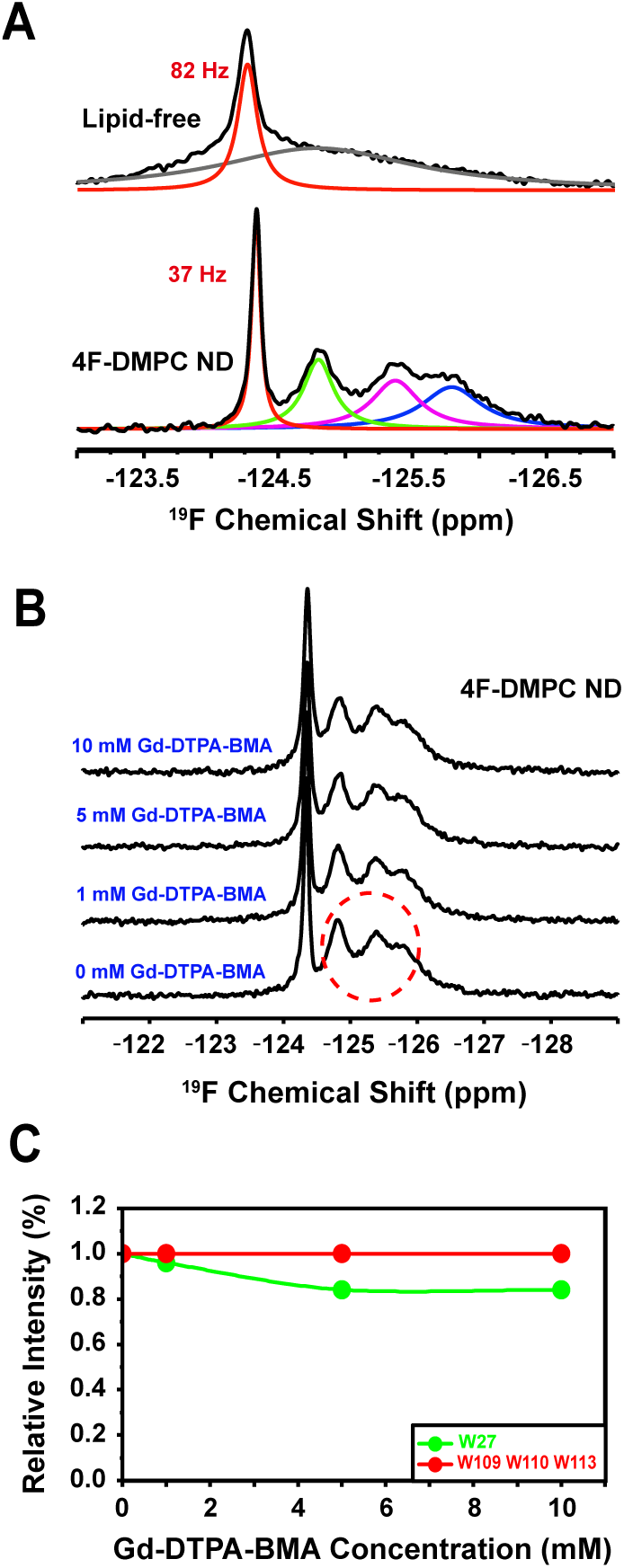
^19^F NMR experiments on nanodiscs containing ^19^F-cytb5. (A) ^19^F NMR spectra of 5F-tryptophan-labeled cytb5 alone in buffer and reconstituted in 4F-DMPC nanodiscs at 318 K. (B) ^19^F NMR spectra of cytb5 reconstituted in 4F-DMPC nanodiscs with varying concentration of Gd-DTPA-BMA. (C) NMR peak intensity observed for tryptophan residues of cytb5 in presence of Gd-DTPA-BMA.

### Effects of different lipid composition on cytb5

Since the structure, folding and topology of a membrane protein could depend on the lipid composition of membrane mimetics used, we investigated the effects of varying the lipid composition on the reconstitution of cytb5. For this purpose, in addition to 4F-DMPC nanodiscs, 4F-POPC, 4F-POPC/PS *(8:2 molar ratio)* and 4F-POPS nanodiscs were prepared as described above. The diameters of these nanodiscs were measured to be ~8.4 nm for 4F-POPC, ~8.0 nm for 4F-POPC/PS *(8:2 molar ratio)*, ~6.2 nm for 4F-POPS (Figure 4). We reconstituted 5FW-labeled cytb5 into these nanodiscs as described above. SEC and DLS profiles and ^19^F NMR spectra indicate that cytb5 is reconstituted into these nanodiscs that vary in the lipid composition (Figure 4).

**Figure 4.**
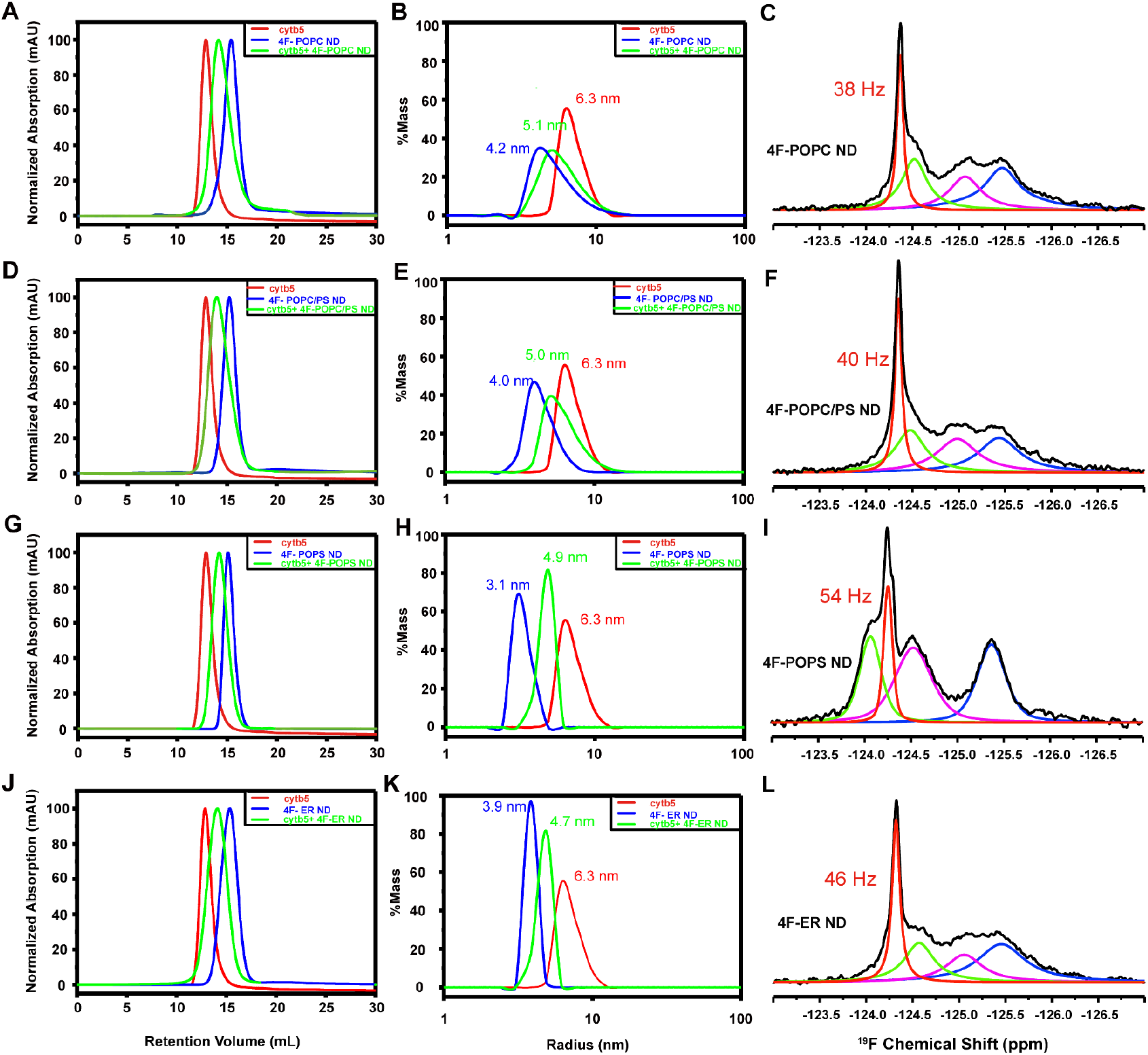
Characterization of 4F-peptide nanodiscs containing ^19^F-cytb5. Reconstitution of cytb5 in 4F-POPC nanodiscs, 4F-POPC/PS nanodiscs, 4F-POPS nanodiscs and 4F-ER nanodiscs. SEC elution profiles (A, D, G, J) and DLS profiles (B, E, H, K) of cytb5 alone in buffer (red), empty nanodiscs (blue), and nanodiscs containing cytb5 (green). Fits of ^19^F NMR spectra of 5F-tryptophan-labeled cytb5 reconstituted in 4F-POPC (C), 4F-POPC/PS (F), 4F-POPS (I), and 4F-ER nanodiscs(L) at 310 K.

^19^F NMR spectra of cytb5 reconstituted in 4F-POPC, 4F-POPC/PS *(8:2 molar ratio)*, or 4F-POPS nanodiscs were also acquired at various temperatures, and the spectra are shown in Figure 2. All of these spectra exhibit a narrow peak for W27 with a line width of 57 Hz in 4F-POPC, 55 Hz in 4F-POPC/PS, and 65 Hz in 4F-POPS. With the increasing temperature, besides the narrow peak from W27, three ^19^F NMR peaks from Trp residues in the transmembrane domain are also observed in all three types of nanodiscs at high temperatures (Figure 2). The presence of zwitterionic lipids like DMPC or POPC in nanodiscs resulted in the same ^19^F NMR peak positions indicating the uncharged lipids have a negligible effect on the soluble domains of cytb5. This is in complete agreement with previous NMR studies on membrane-bound cytb5 that reported a fast time scale of motion for residues from the soluble domain with negligible interaction with zwitterionic lipid membrane. On the other hand, with a gradual increase in the concentration of negatively charged POPS lipids, a low-field shift was observed for the ^19^F peak from W27 (Figures 2 and 4). However, for the residues from the transmembrane domain that exhibited broader lines as compared to that from W27, the increased chain length and the presence of an unsaturated bond in the neutral POPC lipids induced moderate chemical shifts as compared to that observed from DMPC nanodiscs. On the other hand, with the increasing concentration of negatively charged POPS, ^19^F spectra appeared to be significantly different from that observed from zwitterionic PC only lipids containing nanodiscs. The presence of 100 % POPS resulted in a maximum change in the chemical shift (Figure 2).

Cytb5 is primarily associated with the endoplasmic reticulum of eukaryotic cells. Therefore, biomimetic 4F-ER nanodiscs were developed to mimic the natural lipid membrane environment for cytb5. The diameter of the 4F-ER nanodiscs was determined to be ~8 nm (Figure 4). The SEC profiles and ^19^F NMR spectra (Figure 4) confirmed the successful reconstitution of cytb5 in 4F-ER nanodiscs.

### Formation of cytb5-CYP2B4 complex in nanodiscs

Upon the addition of an unlabeled full-length CYP2B4 to nanodiscs containing cytb5, no obvious chemical shift perturbation was observed for the ^19^F peak from W27, but the intensity decreased significantly as compared to that observed for cytb5 alone (Figure 5), suggesting the formation of a productive cytb5-CYP2B4 complex in the nanodiscs. The complex formation was also confirmed by performing 2D TROSY-HSQC NMR experiments as reported previously.[20, 22] For the residues from the transmembrane domain, a small chemical shift perturbation was observed with a significant reduction in the signal intensity (Figure 5). The addition of BHT to the cytb5-CYP2B4 further broadened the ^19^F peak from W27 to beyond the detection limit, suggesting a tighter binding between cytb5 and CYP2B4 in presence of BHT. These results are consistent with our previous findings in DMPC/DHPC isotropic bicelles[20], and 22A-nanodiscs[21] by using ^15^N-cytb5 and CYP2B4.

**Figure 5.**
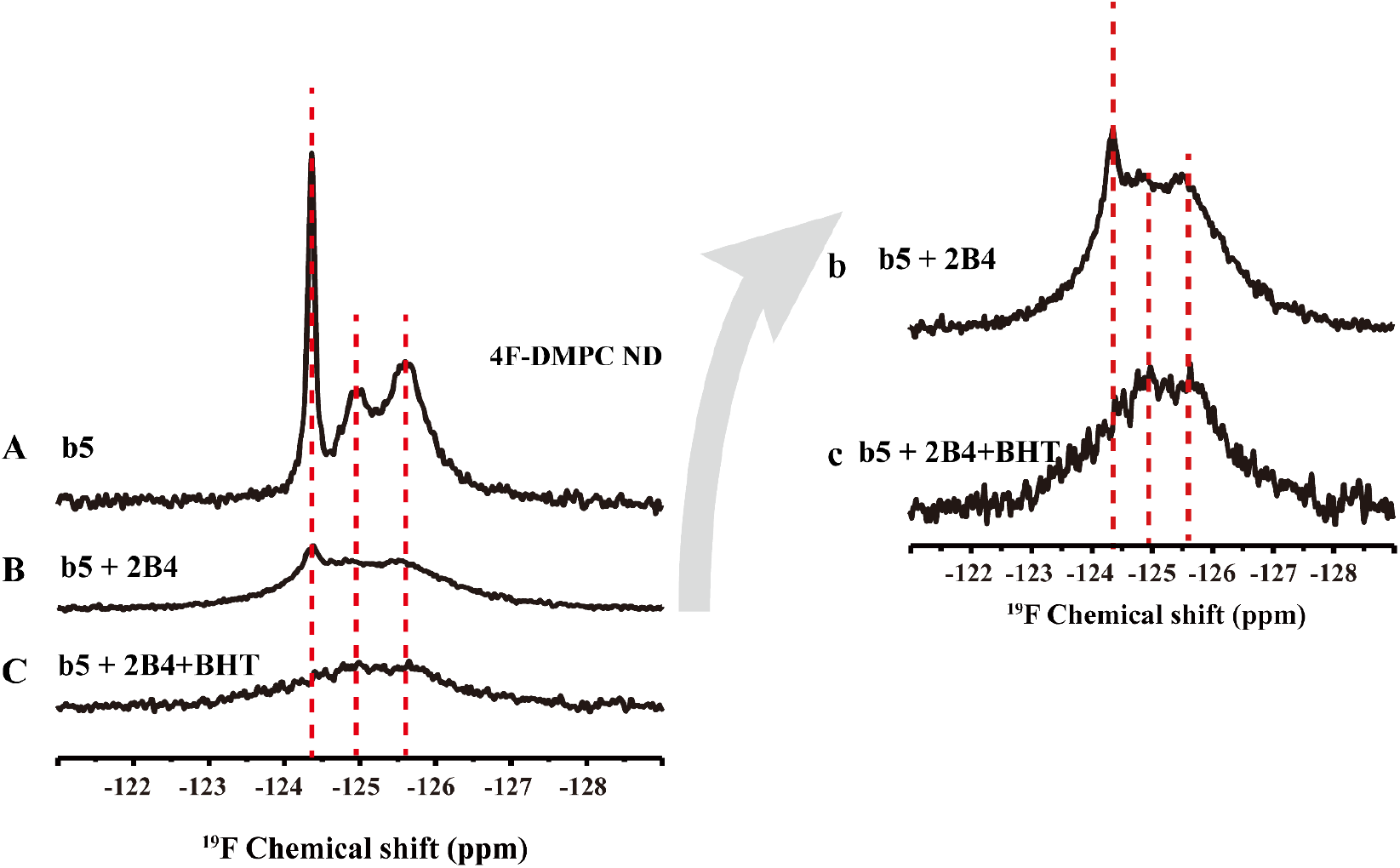
^19^F NMR revealing cytb5-cytP450 interactions in nanodiscs. ^19^F NMR spectra of 5F-tryptophan-labeled cytb5 alone in 4F-DMPC nanodiscs (A), cytb5 in complex with P450 2B4 (B), and cytb5-P450 in complex in presence of BHT at 298K. (b) and (c) are the magnification of (B) and (C), respectively.

## Conclusions

In this study, wild-type cytb5 labeled with 5-fluorotryptophan (5FW) was successfully reconstituted in peptide-based nanodiscs with a varying lipid composition including 4F-DMPC, 4F-POPC, 4F-POPC/PS, 4F-POPS and 4F-ER nanodiscs. Simple ^19^F NMR spectra obtained from nanodiscs composed of DMPC, POPC, POPC/PS, or ER lipids and cytb5 at different temperatures demonstrated the feasibility of resolving the spectral lines from the transmembrane domains of the protein. A significant change in the chemical shift was observed in presence of a negative charged lipid POPS. Interaction of cytb5 with CYP2B4 resulted in broadening of all the four 19F spectral lines and further broadening was observed in presence of BHT indicating a tight complex formation. These results demonstrate the feasibility of ^19^F NMR experiments on nanodiscs containing single-pass transmembrane proteins.

## ACKNOWLEDGEMENTS

This study was supported by NIH (GM084018 to A.R.).

